# Divergent PAM Specificity of a Highly-Similar SpCas9 Ortholog

**DOI:** 10.1101/258939

**Authors:** Pranam Chatterjee, Noah Jakimo, Joseph M. Jacobson

## Abstract

RNA-guided DNA endonucleases of the CRISPR-Cas system are widely used for genome engineering and thus have numerous applications in a wide variety of fields. The range of sequences that CRISPR endonucleases can recognize, however, is constrained by the need for a specific protospacer adjacent motif (PAM) flanking the target site. In this study, we demonstrate the natural PAM plasticity of a highly-similar, yet previously uncharacterized, Cas9 from *Streptococcus canis* (ScCas9) through rational manipulation of distinguishing motif insertions. To this end, we report a divergent affinity to 5’-NNGT-3’ PAM sequences and demonstrate the editing capabilities of the ortholog in both bacterial and human cells. Finally, we build an automated bioinformatics pipeline, the Search for PAMs by ALignment Of Targets (SPAMALOT), which further explores the microbial PAM diversity of otherwise-overlooked *Streptococcus* Cas9 orthologs. Our results establish that ScCas9 can be utilized both as an alternative genome editing tool and as a functional platform to discover novel *Streptococcus* PAM specificities.

## Introduction

RNA-guided endonucleases of the CRISPR-Cas system, such as Cas9 (1) and Cpf1 (also known as Cas12a) (2), have proven to be versatile tools for genome editing and regulation (3), which have numerous implications in medicine, agriculture, bioenergy, food security, nanotechnology, and beyond (4). The range of targetable sequences is limited, however, by the need for a specific protospacer adjacent motif (PAM), which is determined by DNA-protein interactions, to immediately follow the DNA sequence specified by the single guide RNA (sgRNA) (1, 5–8). For example, the most widely used variant, *Streptococcus pyogenes* Cas9 (SpCas9), requires an 5’-NGG-3’ motif downstream of its RNA-programmed DNA target (1,4–8). To relax this constraint, additional Cas9 and Cpf1 variants with distinct PAM requirements have been either discovered (9–13) or engineered (12–15) to diversify the range of targetable DNA sequences. In total, these studies have provided only a handful of CRISPR effectors with minimal PAM requirements that enable wide targeting capabilities.

To help augment this list, we characterize an orthologous Cas9 protein from *Streptococcus canis*, ScCas9 (UniProt I7QXF2), possessing 89.2% sequence similarity to SpCas9. We find that despite such homology, ScCas9 prefers a non-canonical 5’-NNGT-3’ PAM. To explain this divergence, we identify two significant insertions within its open reading frame (ORF) that differentiate ScCas9 from SpCas9 and contribute to its PAM-recognition flexibility. We show that ScCas9 can efficiently edit genomic DNA in mammalian cells, and construct a bioinformatics pipeline to explore the PAM specificities of other *Streptococcus* orthologs. Finally, we investigate possible explanations for PAM divergence between *Streptococcus* orthologs by combining our engineering and bioinformatics approaches.

## Results

### Identification of SpCas9 Homologs

While numerous Cas9 homologs have been sequenced, only a handful of *Streptococcus* orthologs have been characterized or functionally validated. To explore this space, we curated all *Streptococcus* Cas9 protein sequences from UniProt (16), performed global pairwise alignments using the BLOSUM62 scoring matrix (17), and calculated percent sequence homology to SpCas9. From them, the Cas9 from *Streptococcus canis* (ScCas9) stood out, not only due to its remarkable sequence homology (89.2%) to SpCas9, but also because of a positive-charged insertion of 10 amino acids within the highly-conserved REC3 domain, in positions 367–376 Figure 1A. Exploiting both of these properties, we modeled the insertion within the corresponding domain of PDB 4OO8 (18) and, when viewed in PyMol, noticed that it formed a “loop”-like structure, of which several of its positive-charged residues come in close proximity with the target DNA near the PAM Figure 1B. We further identified an additional insertion of two amino acids (KQ) immediately upstream of the two critical arginine residues necessary for PAM binding (19), in positions 1337–1338 Figure 1A. We thus hypothesized that these insertions may affect the PAM specificity of this enzyme. To support this prediction, we computationally characterized the PAM for ScCas9, by first mapping spacer sequences from the Cas9-associated type II CRISPR loci in the *Streptococcus canis* genome (20) to viral and plasmid genomes using BLAST (21), extracting the sequences 3’ to the mapped protospacers, and subsequently generating a WebLogo (22) representation of the aligned PAM sequences. Our analysis suggested an 5’-NNGTT-3’ PAM Figure 1C. Intrigued by these novel motifs and motivated by its predicted, divergent PAM, we selected ScCas9 as a candidate for further PAM characterization and engineering.

**Figure 1.**
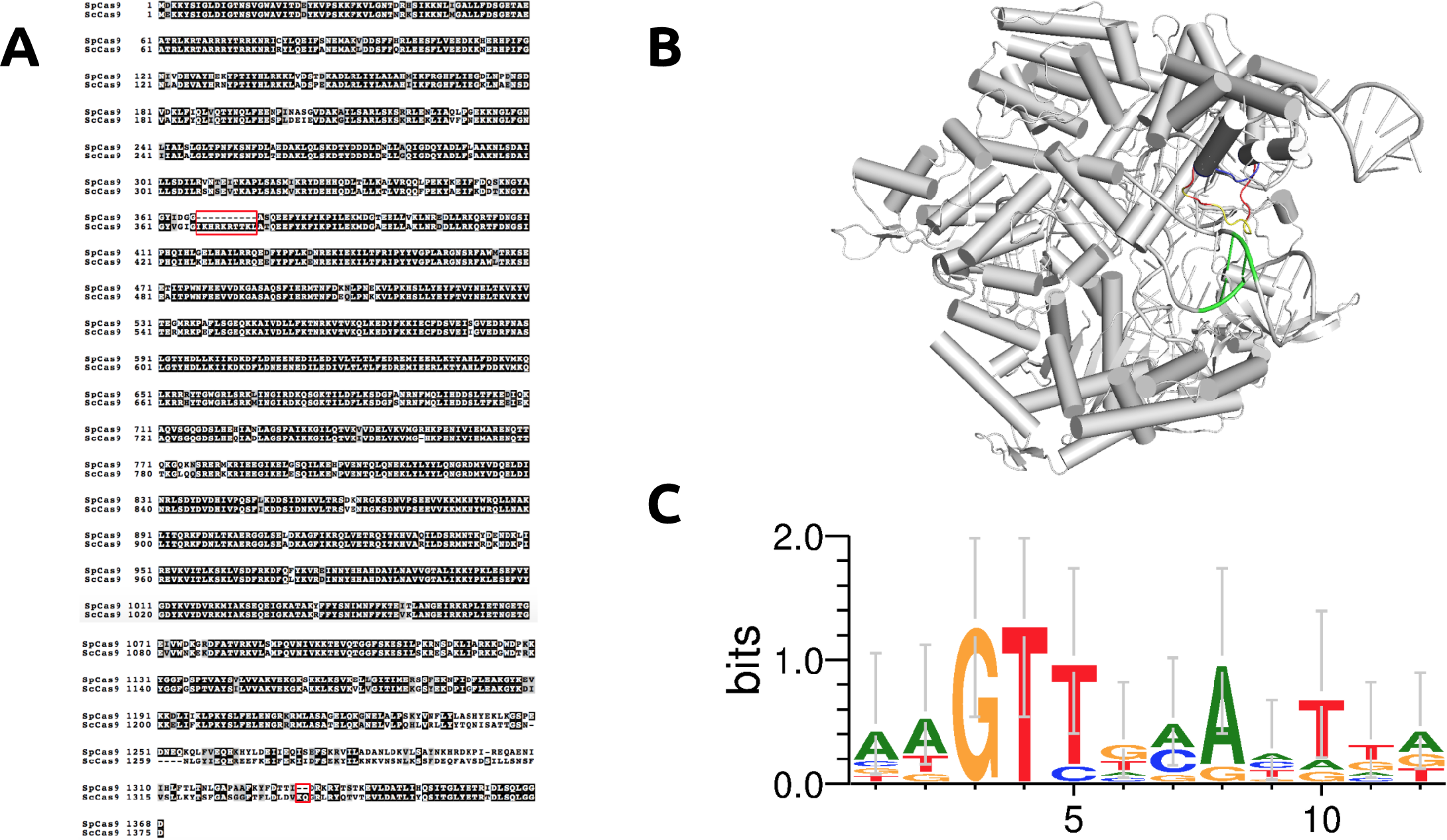
Elucidation of ScCas9. (A) Global pairwise sequence alignment of SpCas9 and ScCas9. Despite sharing 89.2% sequence homology to SpCas9, ScCas9 contains two notable insertions, one positive-charged insertion in the REC domain (367–376) and another KQ insertion in the PAM-interacting domain (1337–1338), as indicated. B) Insertion of novel REC motif into PDB 4OO8 (18). The 367–376 insertion demonstrates a loop-like structure (red). Several of its positive-charged residues (yellow) come in close proximity to the target DNA near the PAM (green). C) WebLogo (22) for sequences found at the 3’ end of protospacer targets identified in plasmid and viral genomes using Type II spacer sequences within *Streptococcus canis* as BLAST (20) queries.

### Determination of PAM Sequences Recognized by ScCas9

Due to the the relatively low number of protospacer targets, we validated the PAM binding sequence of ScCas9 utilizing an existent positive selection bacterial screen based on GFP expression conditioned on PAM binding, termed PAM-SCANR (23). A plasmid library containing the target sequence followed by a randomized 5’-CNNNNC-3’ PAM sequence was bound by a nuclease-deficient ScCas9 (and dSpCas9 as a control) and an sgRNA both specific to the target sequence and general for SpCas9 and ScCas9, allowing for the repression of *lacI* and expression of GFP. Plasmid DNA from FACS-sorted GFP-positive cells and pre-sorted cells were extracted and amplified, and enriched PAM sequences were identified by Sanger sequencing. Our results provide initial evidence that ScCas9 can bind to a more minimal 5’-NNGT-3’ PAM, distinct to that of SpCas9’s 5’-NGG-3’ Figure 2A.

**Figure 2.**
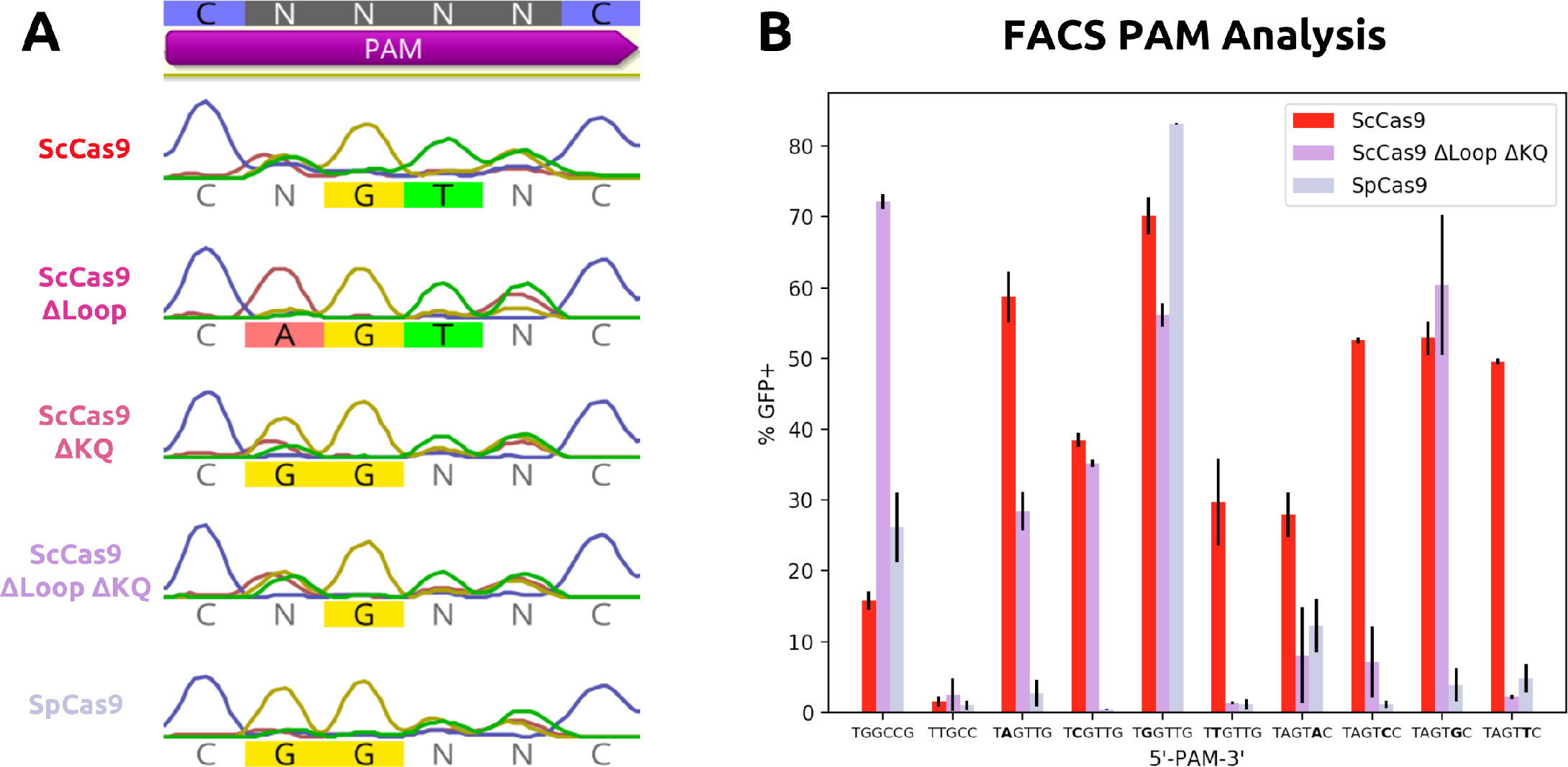
PAM Determination of Engineered ScCas9 Variants. A) PAM binding enrichment on a 5’-CNNNNC-3’ PAM library. We implemented PAM-SCANR (23) and observed divergent PAM specificity of ScCas9 (5’-NNGT-3’) relative to SpCas9 after Sanger sequencing selected cell populations. Removal of the loop (∆367–376) generates a variant that prefers a longer 5’-NAGT-3’ sequence, while removal of the KQ insert (∆1337–1338) reverts the PAM preference to a more SpCas9-like 5’-NGG-3’ sequence. Removal of both motifs predicts a potential 5’-NNG-3’, with minimal preference for T at position 4. B) Examination of PAM preference for ScCas9. For individual PAMs, we varied a single position (2 and 5) to test each possible base. Our results confirm ScCas9 activity on 5’-NNGTN-3’ via FACS analysis for GFP expression after electroporation and overnight incubation. ScCas9 ∆Loop ∆KQ demonstrates intermediate PAM specificity with activity on 5’-NGG-3’ but also on certain 5’-NNGTN-3’ PAM sequences.

We hypothesized that the previously described insertions may contribute to this flexibility, and thus engineered ScCas9 to remove either insertion or both, and subjected these variants to the same screen. Only removing the loop (ScCas9 ∆367-376 or ScCas9 ∆Loop) extended the PAM of ScCas9 to 5’-NAGT-3’, while only removing the KQ insertion (ScCas9 ∆1337-1338 or ScCas9 ∆KQ), reverted the PAM specificity to a more 5’-NGG-3’-like PAM with minimal requirements for T at position 4 Figure 2A. Finally, the most SpCas9-like variant, where both insertions are removed (ScCas9 ∆367-376 ∆1337–1338 or ScCas9 ∆Loop ∆KQ), showed a strong preference for G in position 3 while also exhibiting reduced affinity for T at position 4 Figure 2A.

To confirm the results of the library assay, we decided to elucidate the minimal PAM requirements of ScCas9 and ScCas9 ∆Loop ∆KQ by utilizing fixed PAM sequences. We replaced the PAM library with individual PAM sequences, which were varied at positions 2 and 5 to test each possible base. Our results demonstrate that while ScCas9 exhibits a clear 5’-NNGTN-3’ preference, with activity for all bases at both positions, ScCas9 ∆Loop ΔKQ demonstrates significant binding at 5’-NGG-3’ PAM sequences and at some, but not all, 5’-NNGTN-3’ motifs, indicating an intermediate PAM specificity between that of SpCas9 and ScCas9 Figure 2B.

### Genome Editing by ScCas9 Variants in Human Cells

We assessed the ability of ScCas9 and ScCas9 ∆Loop ∆KQ to edit mammalian genomes by co-transfecting HEK293T cells with plasmids expressing these variants along with sgRNAs directed to a native genomic locus (VEGFA) with varying PAM sequences. We first tested editing efficiency at a site containing an overlapping PAM (5’-GGGT-3’). After 48 hours post-transfection, mutation rates detected by the T7E1 assay demonstrated comparable editing activities of SpCas9, ScCas9, and ScCas9 ∆Loop ∆KQ (Figure 3). Additionally, we constructed sgRNAs to sites with various non-overlapping 5’-NNGT-3’ PAM sequences. Other than at the well-described weakly-preferred 5’-NAG-3’ PAM sequence (24, SpCas9’s cleavage activity was abrogated to background levels for other non-overlapping sequences (Figure 3). Alternatively, ScCas9 maintained detectable activity (Figure 3). Consistent with the bacterial data, ScCas9 ∆Loop ∆KQ was able to cleave at most, but not all, non-overlapping 5’-NNGT-3’ sites (Figure 3). Surprisingly, ScCas9 and ScCas9 ∆Loop ∆KQ also exhibited significant cleavage activity at a 5’-GTGAG-3’ PAM (Figure 3), potentially indicating broader PAM specificity than previously determined. Overall, these results verify that ScCas9 can serve as an effective alternative to SpCas9 for genome editing in mammalian cells, both at overlapping 5’-NGGT-3’ and non-overlapping 5’-NNGT-3’ PAM sequences, while also confirming ScCas9 ∆Loop ∆KQ’s intermediate PAM specificity.

**Figure 3.**
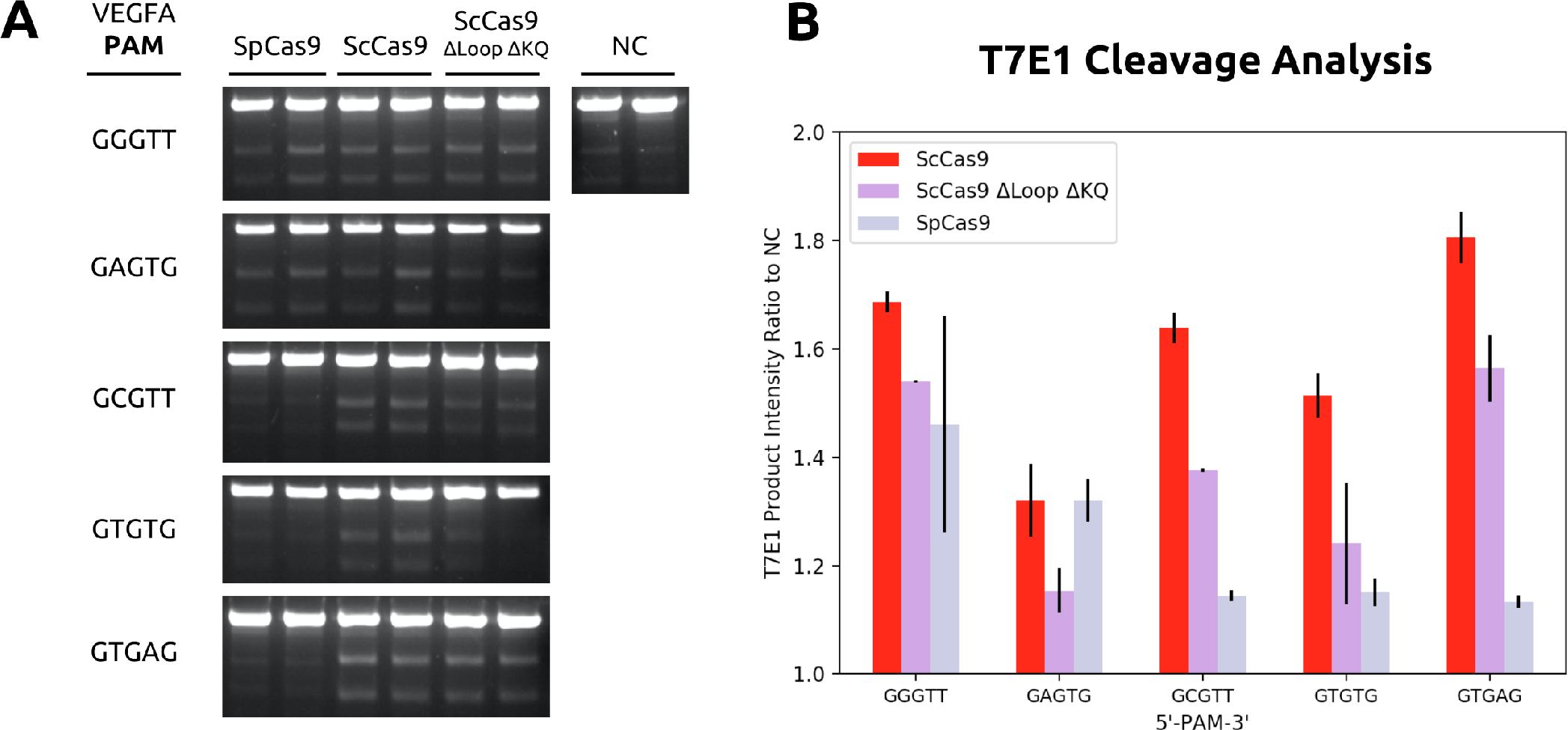
Genome-Editing Activity of ScCas9 in HEK293T Cells. A) T7E1 analysis of indels produced at VEGFA loci with indicated PAM sequences. The Cas9 used is indicated above each lane. B) Quantitative product analysis of T7E1 products. Unprocessed gel images were quantified by line scan analysis using Fiji (34), and represented as ratios to the negative control (NC). Significant gene editing activity was detected for SpCas9, ScCas9, and ScCas9 ∆Loop ∆KQ at an overlapping 5’-NGGT-3’ PAM. Furthermore, ScCas9 displays cleavage abilities at various other sites with 5’-NNGT-3’ PAM sequences. ScCas9 ∆Loop ∆KQ cleaves at some, but not all, of these sites. ScCas9 and ScCas9 ∆Loop ∆KQ also exhibits preference for a 5’-GTGAG-3’ PAM. Finally, SpCas9 demonstrated previously-described cleavage activity on an 5’-NAG-3’ PAM *(24).*

### Genus-wide Prediction of Divergent Streptococcus Cas9 PAMs

Demonstrations of efficient genome editing by Cas9 nucleases with distinct PAM specificity from several *Streptococcus* species, including *S. canis*, motivated us to develop a bioinformatics pipeline for discovering additional Cas9 proteins with novel PAM requirements in the *Streptococcus* genus. We call this method the **S**earch for **PAM**s by **AL**ignment **O**f **T**argets (SPAMALOT). Briefly, we mapped a 20 nt portion of spacers flanked by known *Streptococcus* repeat sequences to candidate protospacers that align with no more than two mismatches in phages associated with the genus. We grouped 12 nt protospacer 3’-adjacent sequences from each alignment by genome and CRISPR repeat, and then generated group WebLogos (22) to compute presumed PAM features. Figure 4A shows that resulting WebLogos accurately reflect the known PAM specificities of Cas9 from *S. canis* (this work), *S. pyogenes, S. thermophilus*, and *S. mutans* (7,25,26). We identified a notable diversity in the WebLogo plots derived from various *S. thermophilus* cassettes with common repeat sequences (Figure 4B), each of which could originate from any other such *S. thermophilus* WebLogo upon subtle specificity changes that traverse intermediate WebLogos among them. We observe a similar relationship between two *S. oralis* WebLogos that also share this repeat, as well as unique putative PAM specificities associated with CRISPR cassettes containing *S. mutans*-like repeats from the *S.oralis, S. equinis*, and *S. pseudopneumoniae* genomes (Figure 4C).

**Figure 4.**
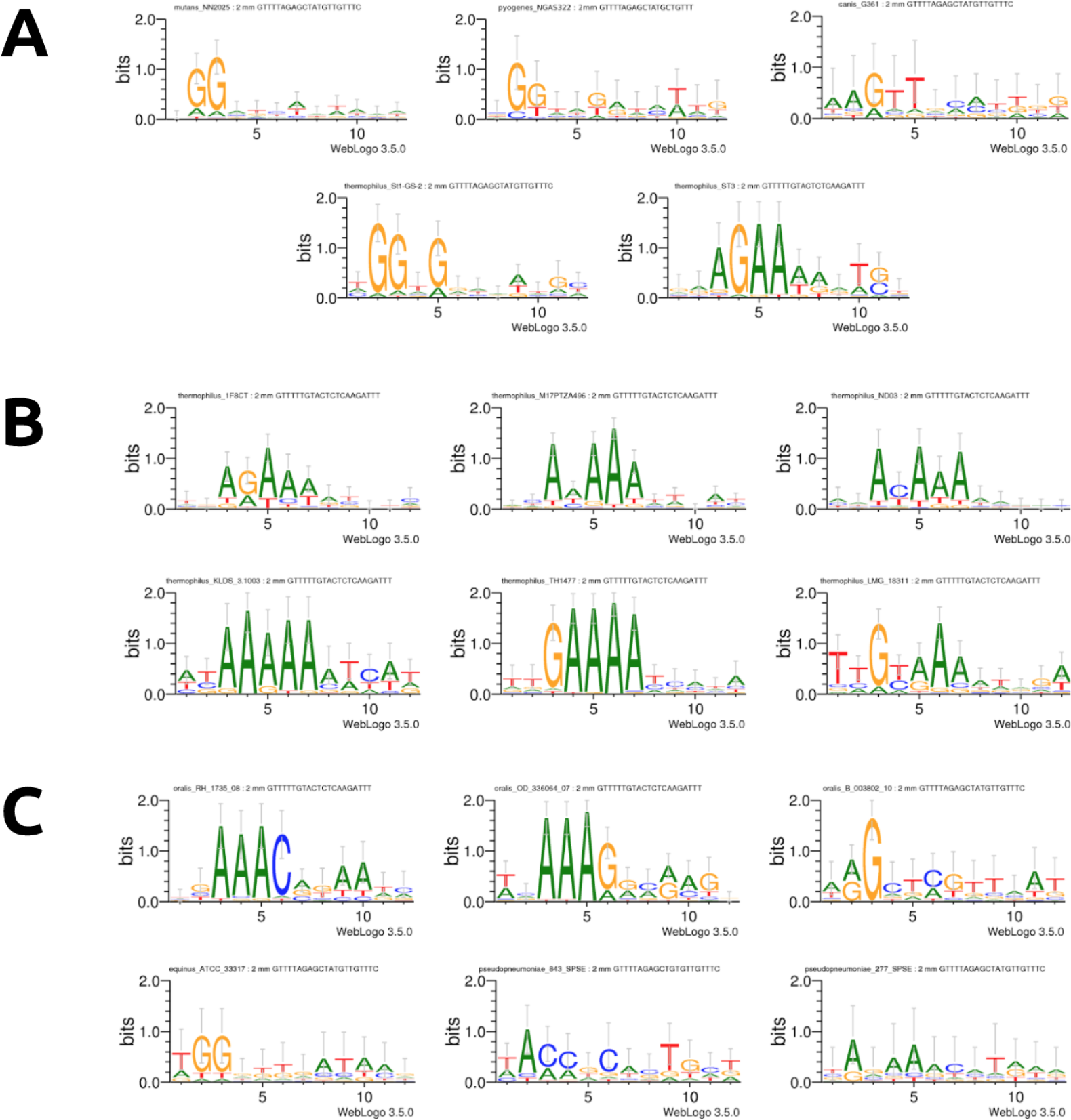
SPAMALOT PAM Predictions for Streptococcus Cas9 Orthologs. Spacer sequences found within the Type II CRISPR cassettes associated with Cas9 ORFs from specified *Streptococcus* genomes were aligned to *Streptococcus phage* genomes to generate spacer-protospacer mappings. WebLogos (22), labeled with the relevant species, genome, and CRISPR repeat, were generated for sequences found at the 3’ end of candidate protospacer targets with no more than two mismatches (2mm). A) PAM predictions for experimentally validated Cas9 PAM sequences in previous studies. SPAMALOT correctly predicts the PAM of experimentally characterized Cas9 enzymes from the specified *S.mutans, S. pyogenes, S. canis,* and *S. thermophilus* genomes (7,25,26). B) Novel PAM predictions of alternate *S. thermophilus* Cas9 orthologs with putative divergent specificities. C) Novel PAM predictions of uncharacterized *Streptococcus* orthologs with distinct specificities.

### Investigation of Sequence Conservation Between S. canis and Other Streptococcus Cas9 Orthologs

To further investigate the distinguishing motif insertions in ScCas9, we inserted the loop (SpCas9 ::Loop), the KQ motif (SpCas9 ::KQ), or both (SpCas9 ::Loop ::KQ) into the SpCas9 ORF and analyzed binding on a 5’-CNNNNC-3’ library. Of these variants, only SpCas9 ::KQ showed target binding affinity in the PAM-SCANR assay. Sequencing on enriched GFP-expressing cells demonstrated an unaffected preference for 5’-NGG-3’ Figure 5A. FACS analysis on a fixed 5’-TGG-3’ PAM confirmed these binding profiles, with SpCas9 ::KQ yielding half the fraction of GFP-positive cells compared to SpCas9 Figure 5B. This data, in conjunction with the binding profiles of ScCas9 variants (Figure 2), suggests that while these insertions within ScCas9 do distinguish its PAM preference from SpCas9, other sequence features of ScCas9 also contribute to its divergence.

**Figure 5.**
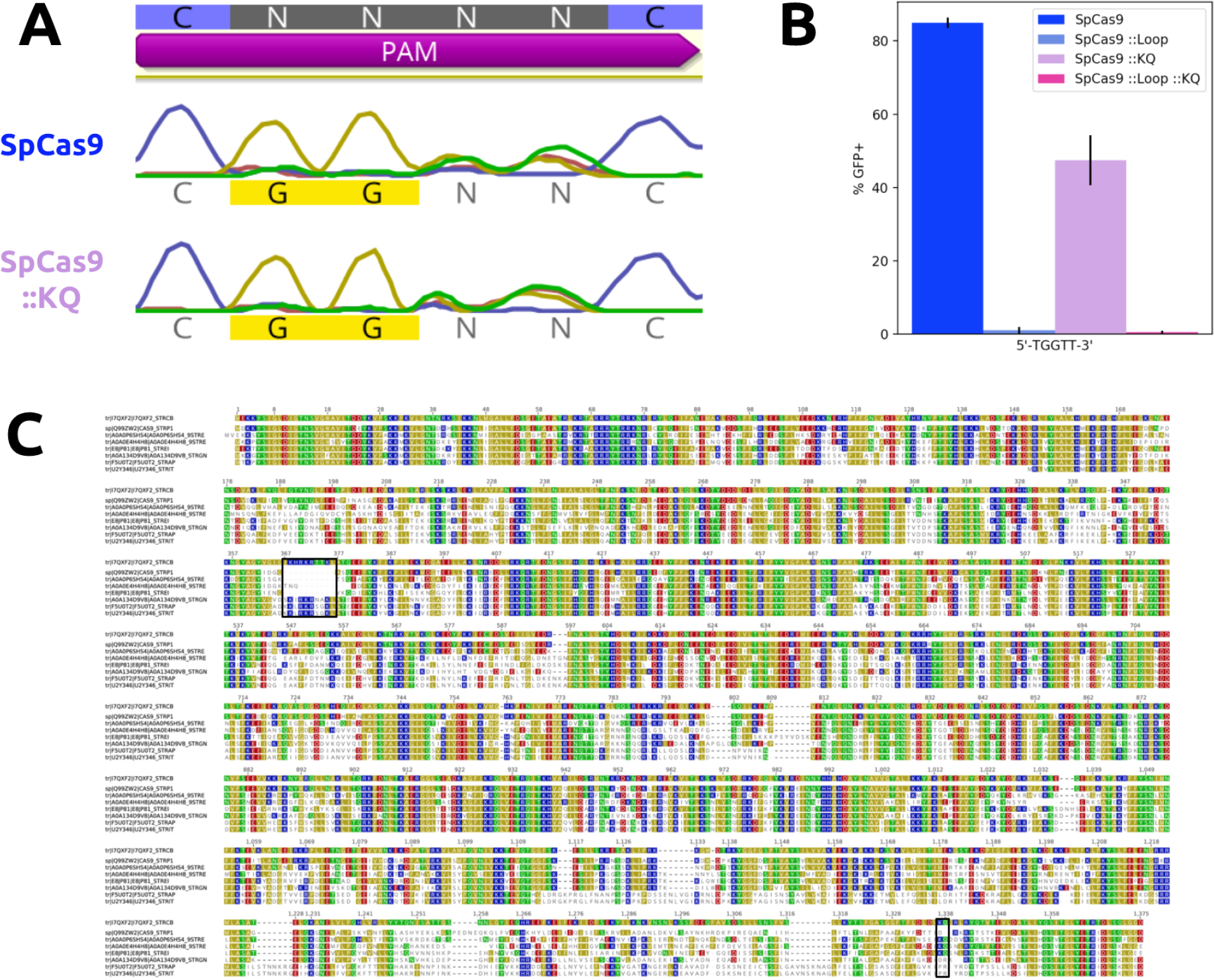
Relationship of ScCas9 to other Streptococcus orthologs. A) PAM binding enrichment on a 5’-CNNNNC-3’ PAM library of ScCas9-like SpCas9 variants. We applied the PAM-SCANR screen (23) to variants of SpCas9 containing either the loop or KQ insertions, or both. While, SpCas9 ::KQ maintained the canonical 5’-NGG-3’ PAM specificity, SpCas9 ::Loop and SpCas9 ::Loop ::KQ failed to demonstrate PAM binding and thus GFP expression. B) Investigation of binding at an 5’-NGG-3’ PAM. Only SpCas9 ::KQ sustains the ability to bind to the canonical SpCas9 PAM, but less effectively than SpCas9. C) Sequence conservation of *Streptococcus* orthologs with ScCas9 as a reference. Each ortholog is referred to by its UniProt ID (16). The loop (367–376) and KQ (1337–1338) insertion alignments are indicated.

*S. canis* has been reported to infect dogs, cats, cows, and humans, and has been implicated as an adjacent evolutionary neighbor of *S. pyogenes*, as evidenced by various phylogenetic analyses (20, 27, 28). In addition to sharing common hosts, we identified *S.canis* CRISPR spacers that map to phage lysogens in *S. pyogenes* genomes, which suggests they are overlapping viral hosts as well. This close evolutionary relationship has manifested itself in the sequence homology of ScCas9 and SpCas9, amongst other orthologous genes, predicted to be a result of lateral gene transfer (LGT) (20, 27, 28). Nonetheless, from the alignment of SpCas9 and ScCas9, the first 1240 positions score with 93.5% similarity and the last 144 positions score with 52.8%. To account for the exceptional divergence in the PAM-interacting domain (PID) at the C-terminus of ScCas9 as well as the positive-charged inserted loop, we focused on alignment of the distinguishing sequences of ScCas9 to other *Streptococcus* Cas9 orthologs (Figure 5C). Notably, the loop motif is present in certain orthologs, such as those from *S. gordonii, S. anginosus*, and *S. intermedius*, while the ScCas9 PID is mostly composed of disjoint sequences from other orthologs, such as those from *S. phocae, S. varani*, and *S. equinis*. Additional LGT events between these orthologs, as opposed to isolated divergence, more likely explain the differences between ScCas9 and SpCas9. Our demonstration that two insertion motifs in ScCas9 alter PAM preferences, yet do not abolish PAM binding when removed, suggests other functional evolutionary intermediates in the formation of effective PAM preferences.

## Discussion

As the growth and development of CRISPR technologies continue, the range of targetable sequences remains limited by the requirement for a PAM sequence flanking a given target site. While significant discovery and engineering efforts have been undertaken to expand this range (9–15), there are still only a handful of CRISPR endonucleases with minimal specificity requirements. Here, we have developed an analogous platform for genome editing using the Cas9 from *Streptococcus canis*, a highly-similar SpCas9 ortholog.

Established PAM engineering methods, such as random mutagenesis and directed evolution, can only generate substitution mutations in protein coding sequences. An alternative approach consists of rationally inserting or removing motifs with specific properties, which may provide a sequence search space that more common mutagenic techniques cannot directly access. Here, we have demonstrated the efficacy of this method on ScCas9, whose sequence disparities with SpCas9 include two divergent motifs. Engineered variants lacking these motifs exhibit altered PAM specificities in both bacterial and human cells. While we do observe minimal inconsistencies in PAM preference between the two assays, which may be explained by PAM-dependent allosteric changes that drive DNA cleavage (19), the PAM divergence of ScCas9 from SpCas9 remains consistent in all tested contexts.

To date, there are no open-source tools or platforms specifically for the prediction of PAM sequences, though prior studies have conducted internal bioinformatics-based characterizations prior to experimental validation (9–11, 13,25). Here, we have established SPAMALOT as a resource that we plan to share with the community for application to CRISPR cassettes from other genera. Future development will include broadening the scope of candidate targets beyond genus-associated phage to capture additional sequences that could be beneficial targets, such as lysogens in species that host the same phage. We hope that this pipeline can be utilized to more efficiently validate and engineer PAM specificities that expand the targeting range of CRISPR, especially for strictly PAM-constrained technologies such as base editing (29) and homology repair induction (30).

Finally, because ScCas9 does not require any alterations to the sgRNA of SpCas9, and due to its significant sequence homology with SpCas9, we presume that identical modifications from previous studies (31–33) can be made to increase the accuracy and specificity of the endonuclease and its variants, though further characterization is necessary. Overall, we anticipate that the development of a broadened *Streptococcus* Cas9 toolkit with expanded targeting range will enhance the current set of CRISPR technologies.

## Materials and Methods

### Identification of Cas9 Homologs and Generation of Plasmids

The UniProt database (16) was mined for all *Streptococcus* Cas9 protein sequences, which were used as inputs to either the BioPython *pairwise2* module or Geneious to conduct global pairwise alignments with SpCas9, using the BLOSUM62 scoring matrix (17), and subsequently calculate percent homology. The Cas9 from *Streptococcus canis* was codon optimized for *E. Coli*, ordered as gBlocks from Integrated DNA Technologies (IDT), and assembled using Golden Gate Assembly. Engineering of the coding sequence of ScCas9 was conducted using either the Q5 Site-Directed Mutagenesis Kit (NEB) or Gibson Assembly. Plasmid backbones for human expression were purchased from Oxford Genetics, and manipulated to individually insert the ORFs of SpCas9, ScCas9 variants, or the sgRNA targeting sequence using Gibson or Golden Gate Assembly.

### Pam-Scanr Assay

Plasmids for the SpCas9 sgRNA and PAM-SCANR genetic circuit, as well as BW25113 ∆lacI cells, were generously provided by the Beisel Lab. Plasmid libraries containing the target sequence followed by either randomized 4-bp 5’-CNNNNC-3’ or fixed PAM sequences were constructed by conducting site-directed mutagenesis on the PAM-SCANR plasmid. Nuclease-deficient mutations were introduced to the Cas9 variants using Gibson Assembly. The provided BW25113 cells were made electrocompetent using standard glycerol wash and resuspension protocols. The dCas9 and sgRNA plasmids, with resistance to chloromphenicol (Chl) and carbenicillin (Crb) respectively, were co-electroporated into the electrocompetent cells at 2.4 kV, outgrown, and plated on Crb+Chl Luria Broth (LB) agar plates overnight. Individual colonies were picked, grown to ABS600 of 0.6 in Crb+Chl LB liquid media, and made electrocompetent. The PAM-SCANR plasmid, with resistance to kanamycin (Kan), was electroporated into the electrocompetent cells harboring both the dCas9 and sgRNA plasmids, outgrown, and collected in 5 mL Crb+Chl+Kan LB media. Overnight cultures were diluted to an ABS600 of 0.01 and cultured to an OD600 of 0.2. Cultures were analyzed and sorted on a FACSAria machine (Becton Dickinson). Events were gated based on forward scatter and side scatter and fluorescence was measured in the FITC channel, with at least 30,000 gated events for data analysis. Sorted GFP-positive cells were grown to sufficient density, and plasmids from the pre-sorted and sorted populations were then isolated, and the region flanking the nucleotide library was PCR amplified and submitted for Sanger sequencing (Genewiz). Bacteria harboring non-library PAM plasmids were analyzed by FACS analysis following electroporation and overnight incubation, and represented as the percent of GFP-positive cells in the population. Additional details on the PAM-SCANR assay can be found in Leenay, et al. (23).

### Cell Culture and Indel Analysis

HEK293T cells were maintained in DMEM supplemented with 100 units/ml penicillin, 100 mg/ml streptomycin, and 10% fetal bovine serum (FBS). sgRNA plasmid (500 ng) and Cas9 plasmid (500 ng) were transfected into cells (2 × 10^5^/well in a 24-well plate) with Lipofectamine 2000 (Invitrogen) in Opti-MEM (Gibco). After 48 hours post-transfection, genomic DNA was extracted using QuickExtract Solution (Epicentre), and VEGFA loci were amplified by PCR. The T7E1 reaction was conducted according to the manufacturers instructions and equal concentration of products were analyzed on a 2% agarose gel stained with SYBR Safe (Thermo Fisher Scientific). Unprocessed gel image files were analyzed in Fiji (34). The cleaved bands of interest were isolated using the rectangle tool, the area under the corresponding peaks were measured, averaged for each sample, and represented as ratios to the background control.

### SPAMALOT Pipeline

All 11,440 *Streptococcus* bacterial and 53 *Streptococcus* associated phage genomes were downloaded from NCBI. CRISPR repeats catalogued for the genus were downloaded from the CRISPRdb hosted by University of Paris-Sud (35). For each genome, spacers upstream of a specific repeat sequence were collected with a toolchain consisting of the fast and memory-efficient Bowtie 2 alignment (36). Each genome and repeat-type specific collection of spacers were then matched to all phage genomes using the original Bowtie short-sequence alignment tool (37) to identify candidate protospacers with at most one, two, or no mismatches. Unique candidates were input into the WebLogo 3 (22) command line tool for prediction of PAM features.

## Acknowledgments

We are very grateful to Ryan Leenay and Dr. Chase Beisel for sharing PAM-SCANR plasmids and cells, and for providing guidance in implementing the PAM-SCANR protocol. We thank Thrasyvoulos Karydis for critical assistance in modeling PDB domain insertions. We also thank Dr. Ed Boyden for access to cell culture, in addition to Dr. Neil Gershenfeld and Dr. Shuguang Zhang for shared lab equipment.

## Author Contributions

P.C. and N.J. conceived and designed experiments. P.C. and N.J. carried out experiments. P.C. and N.J. conducted data analyses. N.J. and P.C. designed and implemented bioinformatics workflows. P.C. and N.J. wrote the paper. P.C, N.J., and J.M.J. reviewed the paper. J.M.J. supervised the project.

